# Phase Transitions in Mutualistic Communities under Invasion

**DOI:** 10.1101/504605

**Authors:** Samuel R. Bray, Yuhang Fan, Bo Wang

## Abstract

Predicting the outcome of species invasion in ecosystems is a challenge of both theoretical and practical significance. Progress so far has been limited by the intractability of the far-from-equilibrium nature of the transitions that occur during invasion events. Here, we address this limitation by solving for the transition dynamics of a cross-feeding community along an analytically tractable manifold defined by the system carrying capacity. We find that species invasion induces discontinuous transitions in the community composition, resembling phase transitions in physical systems. These sharp transitions are emergent properties of species-resource interactions and relate directly to the extent of niche overlap between invasive and native species. The high susceptibility of community structure to small variations in species phenotype and resource conditions near the phase boundaries can explain empirically observed stochasticity and the emergence of tipping points in ecosystems. Moreover, we demonstrate that these phase transitions can be modulated by environmental variations to construct nonlinear organization of species both in space and time.

## 1. Introduction

Introducing novel species to an ecosystem is a fundamental problem that has many practical applications, including probiotic therapy, collective antibiotic resistance, biodiversity preservation, and ecological engineering [1–4]. Introduction of “invader” species can lead to a variety of outcomes, ranging from native community stabilization to complete community collapse and replacement by invader [5,6]. While empirical studies exist, it remains a challenge to predict invasion outcome based on species’ interaction network topography and invader strategy. Previous theoretical works on community structure have mostly focused on either analyzing the stability of an ecosystem under small perturbations around a certain equilibrium (so-called dynamical stability) [10,11], or defining the range of parameters in which the system remains stable and always returns to a fixed point (so-called structural stability) [5,12,13]. However, these studies have not yet addressed how and when an ecosystem may switch from one stable point to another under strong perturbations. The challenge of species invasion is one obvious perturbation of such kind.

Accurately capturing these transitions requires a model valid throughout the far-from-equilibrium ecosystem state space traversed between multiple fixed points, but classical ecological models primarily rely on pairwise species interactions that are fit to a single, specific, equilibrium environmental condition [14]. On the other hand, building a mechanistic model valid across all regimes of system state space would be intractable due to the high biological complexity of even the simplest ecosystems. Here we reason that resource competition between species constrains the ecosystem’s transition to a manifold defined by the carrying capacity of the environment (Fig. 1A). Other paths between fixed points are less feasible as they would allow excess free resources at the expense of species abundance. This constraint to a resource-limited regime allows us to build an analytically tractable model in which both species and their required environmental resources are explicitly considered [15]. The stability and dynamics of such consumer-resource systems near equilibrium have been extensively characterized both theoretically and experimentally [14], providing a well-defined starting point to solve for their behaviors under invasion.

**Figure 1.**
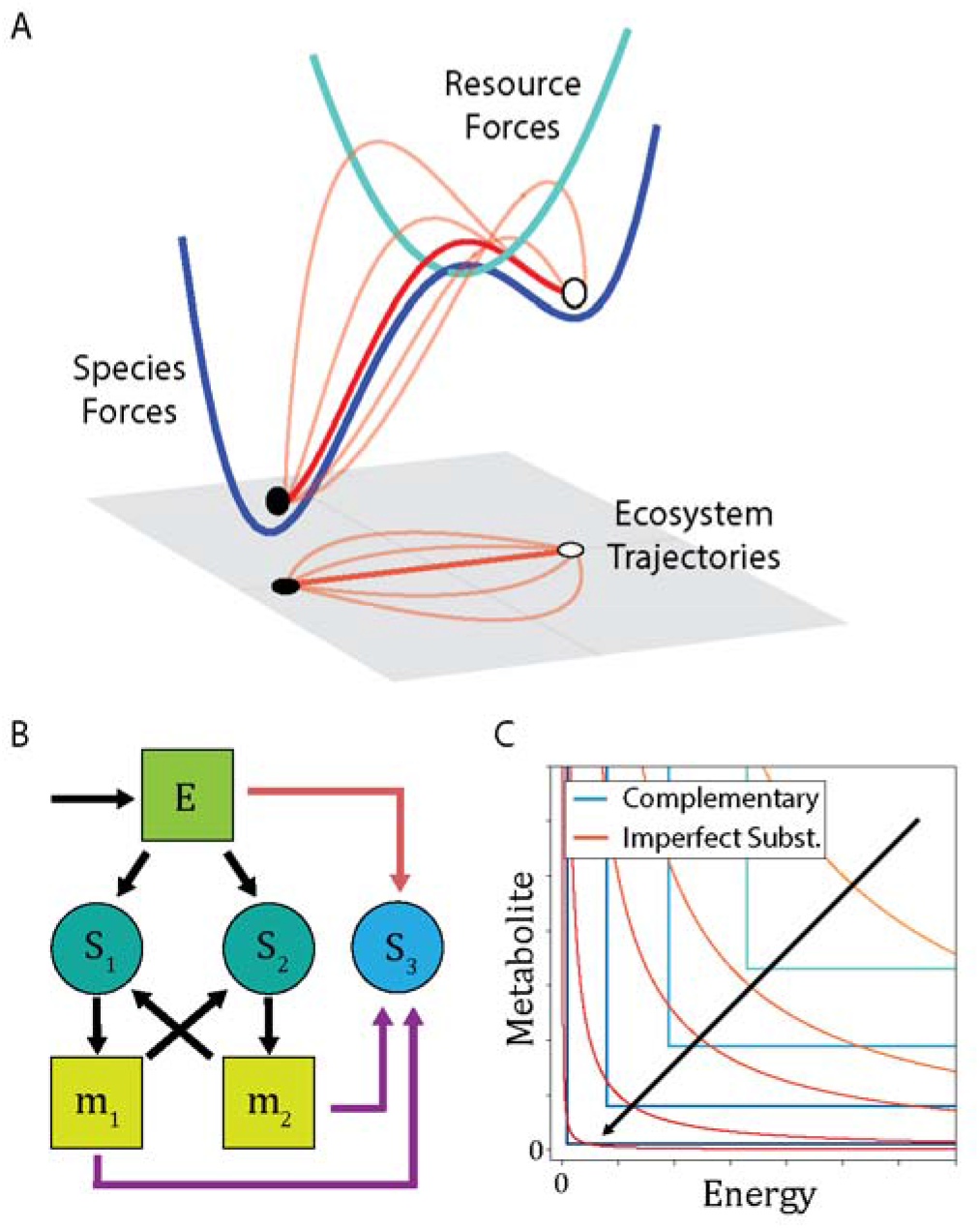
Model mutualistic cross-feeding community under invasion at low-resource limit. (A) Limiting resources constrain the path of state transitions. Species and resource interactions (blue) define fixed points in community structure. Resource availability (green) defines the carrying capacity within which the transition can occur (red). The orange curves represent less feasible paths. (B) In the modeled community, native species, *S*_1_ and *S*_2_, consume a shared energy source *E* and produce unique metabolites *m*_1_ and *m*_2_ required for growth of the complementary species. Invader (*S*_3_) strategies include competition for the energy source (red arrows), parasitic consumption of metabolites (purple arrows), and a mixed strategy that can grow upon both energy and metabolites. Circles: species; squares: resources. (C) Isoclines of growth rates as a function of available resources, energy versus metabolites. Curves of complementary resource condition, i.e., growth only depends on the limiting resource (blue), and imperfectly substitutable resource condition, i.e., growth scales with the product of energy and metabolite abundance (red), converge in the low-resource limit. Arrow indicates the reduction of resource abundance.

The analytical tractability of this model enables us to show that community composition responds to species invasion by undergoing phase transitions, a universal phenomenon in the physical world [16]. These transitions include first-order discontinuous and second-order continuous transitions with discontinuity either in the system state (for first-order transitions) or in the susceptibility of the system to external perturbations (for second-order transitions). The exact outcome can be predicted analytically based on community topography and constraints of environmental resources. The high susceptibility of the system around phase boundaries predicts nonlinear amplifications of stochastic variations during community assembly and species invasion, a phenomenon that has been widely reported in empirical studies [7,9,17,18]. We also discuss the implications of the observed phase transitions in a set of practical problems, including spatial patterns emerging from ecological interactions [19].

## 2. Results

### 2.1 A mechanistic model for cross-feeding community

We consider a metabolic cross-feeding community (Fig. 1B). This type of symbiotic network is abundant in nature, appearing everywhere from auxotrophic bacteria [20,21] to nitrogen fixation in plant root microecosystems [22]. In particular, amino acid cross-feeding has been extensively characterized both theoretically and experimentally [23–26]. Fig. 1B illustrates our two-species model system, defined by state variables representing two species (*S*_1_, *S*_2_), an externally supplied energy source (*E*), and a metabolic product (*m*_l_, *m*_2_) uniquely produced by each species. Each metabolite is required for growth by the other, non-producing species, establishing a syntrophic exchange. The dynamics of each species are controlled by two processes: growth that consumes energy and metabolites at a specific growth rate *µ_i_* and death at the rate *δ*_1_ (Eq. 1). The two metabolites are produced through consumption of energy by their respective species proportional to a production rate *β_i_* and consumed in the other species’ growth (Eq. 2). The energy source is input to the system at a rate *I* and consumed in species growth and metabolite production (Eq. 3).

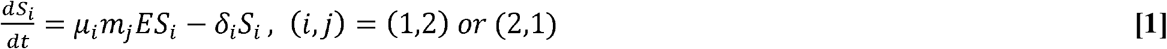

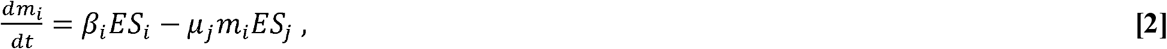

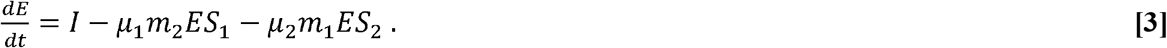

Here we treat the metabolite and energy as “imperfectly substitutable resources” [27], where species growth scales with the product of metabolite and energy abundance. Although this treatment differs from the classical models in which metabolite and energy should be treated as “complementary resources” (i.e., species growth only depends on the limiting resource) [27], it generates a more analytically tractable model and represents a good approximation as it converges onto the classical form when available resources approach zero (Fig. 1C). Since species competition is expected to drive the system towards a resource constrained manifold (Fig. 1A), the low-resource assumptions is valid throughout transitions between fully developed communities. This low-resource limit also allows us to use a linear dependence of growth and resource production on the availability of resources as a first-order approximation of Monod growth [28]. Importantly, our model does not introduce artefactual discontinuities as the system shifts between limiting resources, making it possible to examine the transition dynamics between fixed points of the system.

We analytically identified fixed points by setting all derivatives in Eqs. 1-3 to zero, giving steady state solution (*P*_1_):

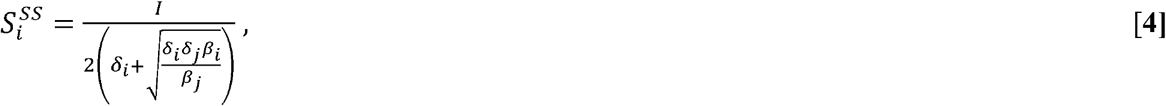

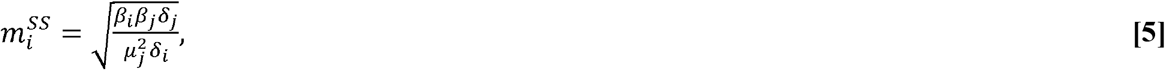

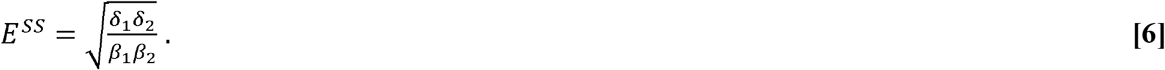

Under constant input of the energy source, the system has only one stable fixed point associated with species coexistence, with the steady state abundance of each species inversely related to the production rate of its unique metabolite. Perturbations to species, resource abundance, and specific growth all maintain species coexistence at the attractor state (see Methods). These results are in accordance with previous experimental and theoretical work demonstrating the stability of cross-feeding communities [23,24].

We next explore how and when the system switches to new fixed points after invasion. We consider three types of invaders: competitors that only use energy (Fig. 1B, red arrows), parasites that only use metabolites produced by the native species (Fig. 1B, purple arrows), and mixed strategy invaders that can use both resources.

### 2.2 Competitive invasion

We first investigated community response to invasion by a competitor. Competitor was added through an additional term for energy consumption by invader growth (−*µ*_3_*ES_3_*) in Eq. 3 and an equation describing invader dynamics:

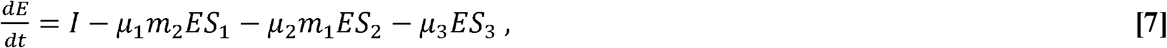

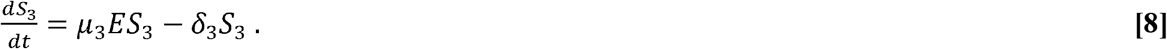

To solve for fixed points, we consider two outcomes separately:

i) Invasion succeeds, which indicates that invader abundance at the fixed point, 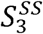, is non-zero. We then consider the case when either native species goes extinct. Given Eqs. 3 and 6, if 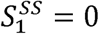, then 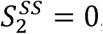, and vice versa. Steady state energy is then:

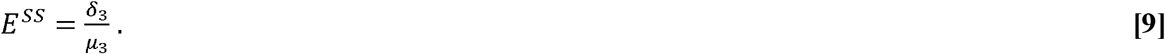

Eq. 7 gives:

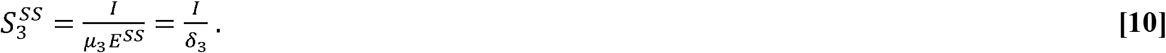

These conditions define a new fixed point, *P*_2_, where the system only contains the invader.

In contrast, if neither native species goes extinct, it can be shown that:

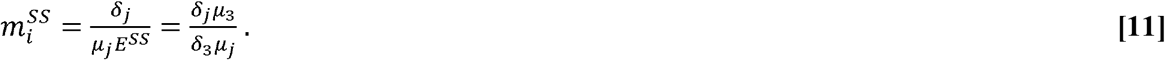

In this condition, Eq. 5 still holds, implying 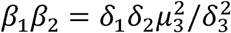, which is not necessarily fulfilled by the model. Therefore, there is no fixed point for three species coexistence.

ii) Invasion fails, which indicates that 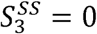. The system degenerates into the form solved for native community only (*P*). The critical condition separating invasion outcomes is determined by the stability of *P*_1_. From Eq. 8, *P*_1_ is stable when *µ*_3_ *E* − *δ*_3_ ≤ 0, leading to a bifurcation in the system at the critical parameter value:

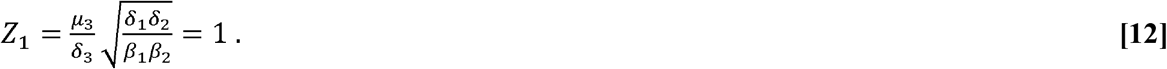

When *Z*_1_ < 1, *P*_1_ is the only stable fixed point. When *Z*_1_ > 1, *P*_1_ becomes unstable to perturbations in invader abundance and *P*_2_ becomes the attractor state (see Methods).

Fig. 2 A shows the phase diagram of the stable steady state community composition in the space of invader specific fitness (*F* = *µ*_3_/*δ*_3_) and community mutualism 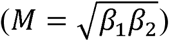. The phase boundary is defined by 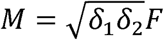. This phase diagram suggests that more rapid conversion of the common energy source into privately shared metabolites strengthens the native community’s cartel of private goods in order to exclude more capable competitors.

**Figure 2.**
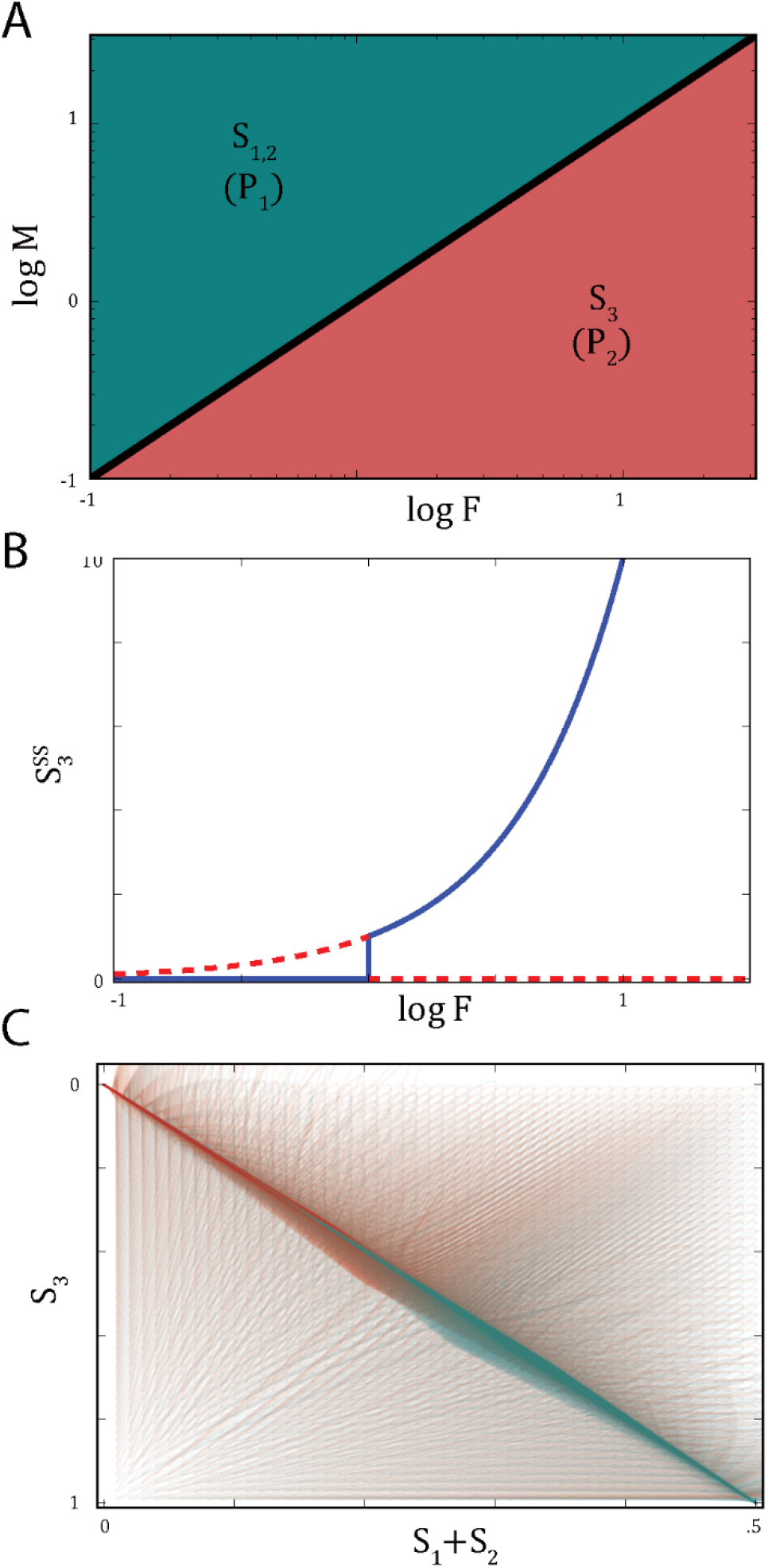
Phase transitions of community composition under competitive invasion. (A) Phase diagram of steady state community composition. Analytically derived phase boundary is marked with black border. (B) Invader abundance plotted against invader fitness for competitive invasion at *M* = 1. Fitness is varied by changing *δ*_3_. Solid line: stable fixed points; dashed lines: unstable points. (C) Time traces of competitive invasion from varied initial conditions rapidly converge to the manifold defined by the carrying capacity in both *P*_1_ (*µ*_3_ = 0.9, cyan) and *P*_2_ (*µ*_3_ = 1.1, red) regimes. Non-varied parameters are all set as 1.

Of particular interest is the change in invader abundance with increasing fitness. In the absence of the native community, there is a single attractor state represented by linear, monotonic increase in invader abundance with increasing fitness (Eq. 10). However, when the complete species network is present, the invader abundance (as well as that of the native species) undergoes a sudden jump representative of a first-order phase transition at the phase boundary between *P*_1_ and *P*_2_ (Fig. 2B). This discontinuity manifests a collective property arising from the network of multispecies interactions. The invader must have a fitness over a critical value to overcome the barrier emerged from the mutualistic interactions within the native community.

The discontinuity at the phase transition serves as an analytical demonstration of the competitive exclusion principle, which states that two species occupying the same niche cannot stably coexist [29]. Here, a niche is defined as a set of resources required for a species to grow. Though the invasive species does not use the exact combination of resources as either native species, it does have the identical *net* resource requirements as the native community as a whole. The sudden extinction of the native community represents its displacement from the energy resource niche.

We then track the dynamics of communities on both sides of the phase boundary across a range of initial conditions to demonstrate that the model is constrained to a resource-limited manifold during the phase transition. The carrying capacity of the system is given as *I* = 2(*δ*_1_*S*_1_ + *δ*_2_*S*_2_) + *δ*_3_*S*_3_. The factor of 2 comes from the cost to convert energy into metabolites. Fig. 2C shows that the community rapidly converges onto the carrying capacity and then moves along this manifold towards its respective fixed point—validating the low-resource assumption in generating the functional form of our model.

### 2.3 Parasitic invasion

With growth only dependent on shared metabolites (Fig. 1B, purple arrows), a parasitic invader exploits the mutualistic interactions of the native community. The original model was modified by an additional term (−*µ*_3_*m*_1_*m*_2_*S*_3_) in Eq. 2 to account for resource consumption by invader growth and an equation describing invader dynamics to give:

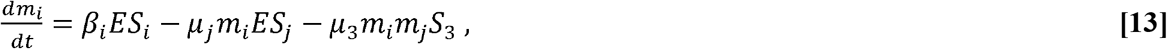

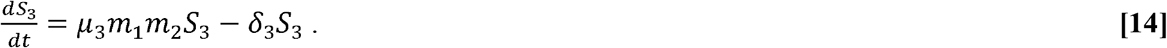

This modification leads to two possible outcomes:

i) Invasion succeeds, which indicates that 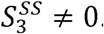. As before, 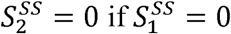, and vice versa. If true, 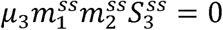 in Eq. 14, leading to 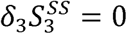, and conflicting with the assumption. Therefore, both 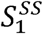 and 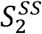 must be non-zero. From Eq. 14 we have:

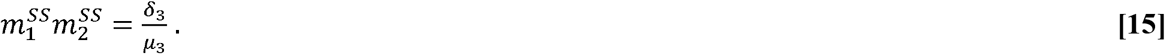

Eqs. 1 and 2 give:

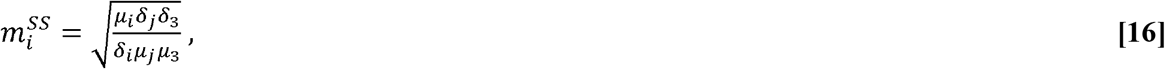

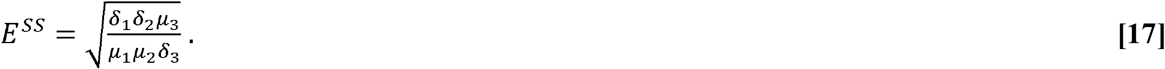

The model then reduces to a system of linear equations for 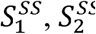 and 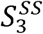, with the solution defining a new fixed point, *P*_3_, where the invader and native community coexist:

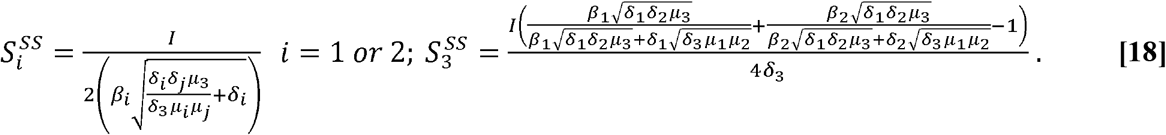

ii) Invasion fails. When 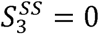, the system degenerates into the form of native community only (*P*_1_). From Eq. 14, the stability *P*_1_ holds when 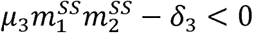, giving a bifurcation point at:

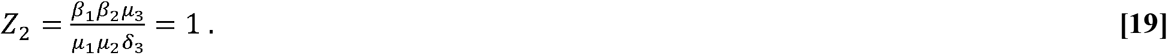

At *Z*_2_ < 1, *P*_1_ is exclusively stable, whereas at *Z*_2_ >1, *P*_3_ is the only stable fixed point (see Methods). Fig. 3 A shows the phase diagram of the two fixed points, again in the space of invader specific fitness and community mutualism, with the phase boundary defined as *M*^2^ = *µ*_1_*µ*_2_*F*^-1^. This phase diagram shows that the native community gains stability with reduced mutualism as the native species divert resources from metabolite production to maximize their own specific growth. This is because, unlike the competitive invasion case, the native species lose exclusive access to shared metabolites and must therefore engage in a growth rate race with the invader to compete for the shared resources.

**Figure 3.**
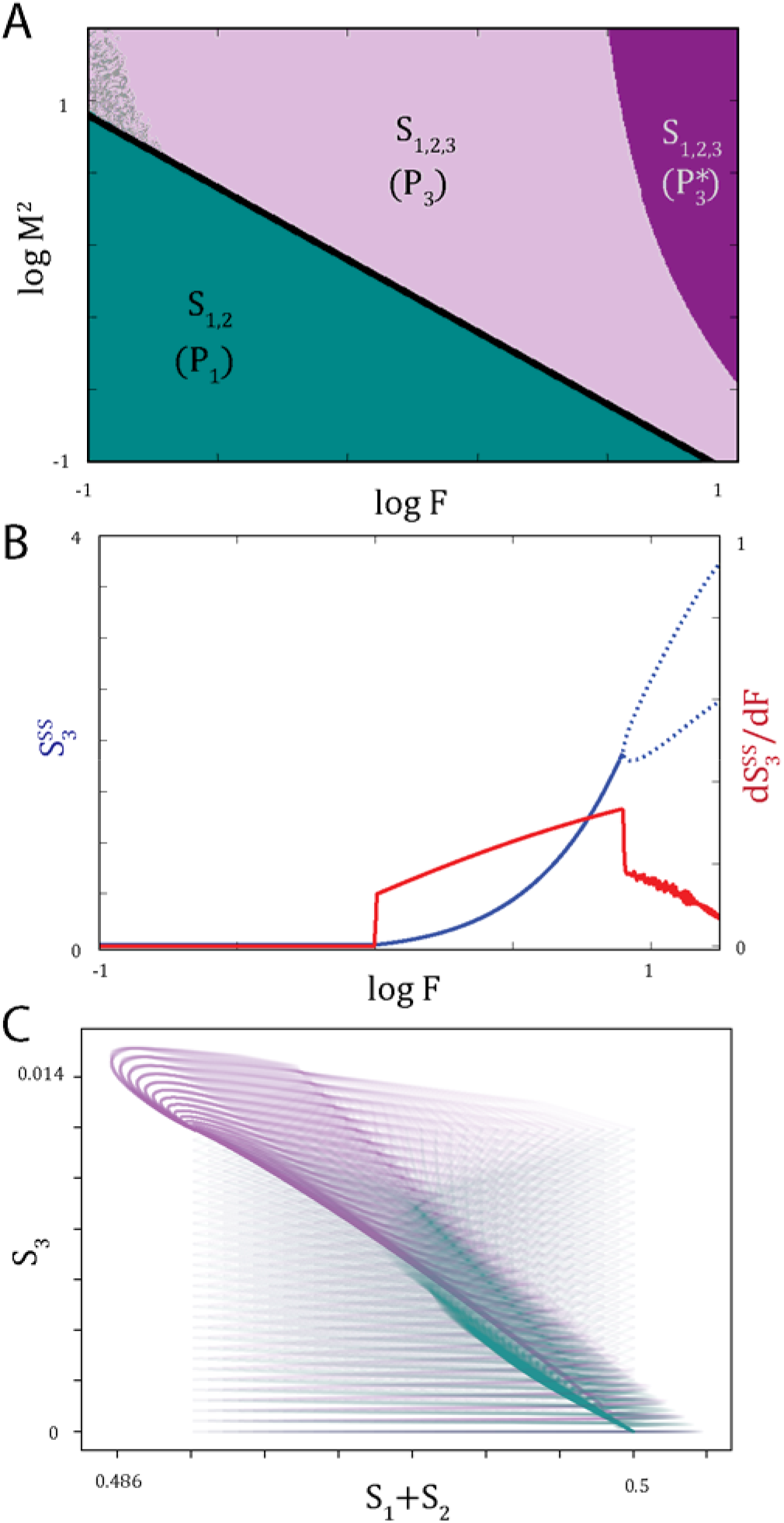
Phase transitions of community composition under parasitic invasion. (A) Phase diagram of steady state community composition. Analytically derived phase boundary is marked with black border. Production rates of metabolite maintain the relation *β*_1_ = *β*_2_. (B) Invader abundance plotted against invader fitness for parasitic invasion at *M* = 1 (blue). Solid line: stable fixed points; dashed line: lower and upper bounds of oscillatory regime around the fixed point corresponding to 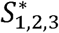 phase in (A). First derivative of abundance (red) shows discontinuity at the phase boundary between fixed points. Invader fitness is varied by changing *δ*_3_. (C) Time traces of parasitic invasion from varied initial conditions converge to the manifold defined by the carrying capacity in both *P*_1_ (*µ*_3_ = 0.99, cyan) and *P*_3_ (*µ*_3_ = 1.01, purple) regimes. Non-varied parameters are all set as 1.

Fig. 3B plots invader abundance across the phase boundary. In contrast to competitive invasion (Fig. 2B), this curve displays a second-order continuous phase transition with the discontinuity only in the derivative of species abundance. In ecology, such transition is also termed as a transcritical bifurcation [30]. Fig. 3C validates that this transition also occurs along the edge of carrying capacity, defined by *I* = 2(*δ*_1_*S*_1_ + *δ*_2_*S*_2_ + *δ*_3_*S*_3_), consistent with the resource-constrained manifold.

Numerical simulation confirmed these two phases and revealed an additional secondary regime wherein a highly mutualistic community coexists with invaders of high specific fitness. Under these conditions, the community enters a stable limit cycle around *P*_3_, creating the new attractor state 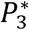 (Fig. 3A-B). This behavior arises from the high load placed on the native community by the parasite. As parasite consumption pushes shared metabolites far below steady state levels it begins to die off, allowing native species to increase in abundance until the metabolite levels recover and allow the parasite to resurge and compete again with the native community.

The different transition order between parasitic and competitive invasions can be interpreted through niche overlap [29]. Because the parasitic invader does not share identical resource requirements with any single or combination of the native species, the species competition only represents partial niche overlap. Increasing invader fitness can thereby continuously expand the distinct niche of the parasite, in contrast to the discontinuity caused by niche displacement in the case of competitive invasion.

### 2.4 Mixed strategy invasion

To integrate both competitive and parasitic invasion strategies in a single invader, we modified Eqs. 2 and 3 for resource consumption as in the previous invasion cases and combined Eqs. 8 and 14 for invader growth:

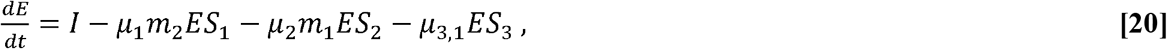

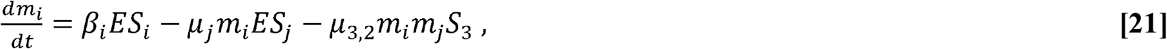

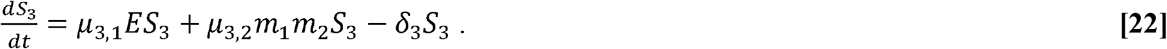

We used numerical analysis to define the phase diagram and identified a triple critical point in community composition (Fig. 4A). With mutualism below the triple point, competition between invader and native community dominates and no coexistence regime exists, creating a first order phase transition between invader rejection and niche replacement. Above the triple point, the increased production of metabolites allows for a coexistence regime with parasitic growth. In this regime, increasing invader fitness first leads to a second order transition as the invader becomes sufficiently competitive to establish the parasitic niche, followed by a first order transition as the invader outcompetes and displaces the native community from the energy resource niche.

**Figure 4.**
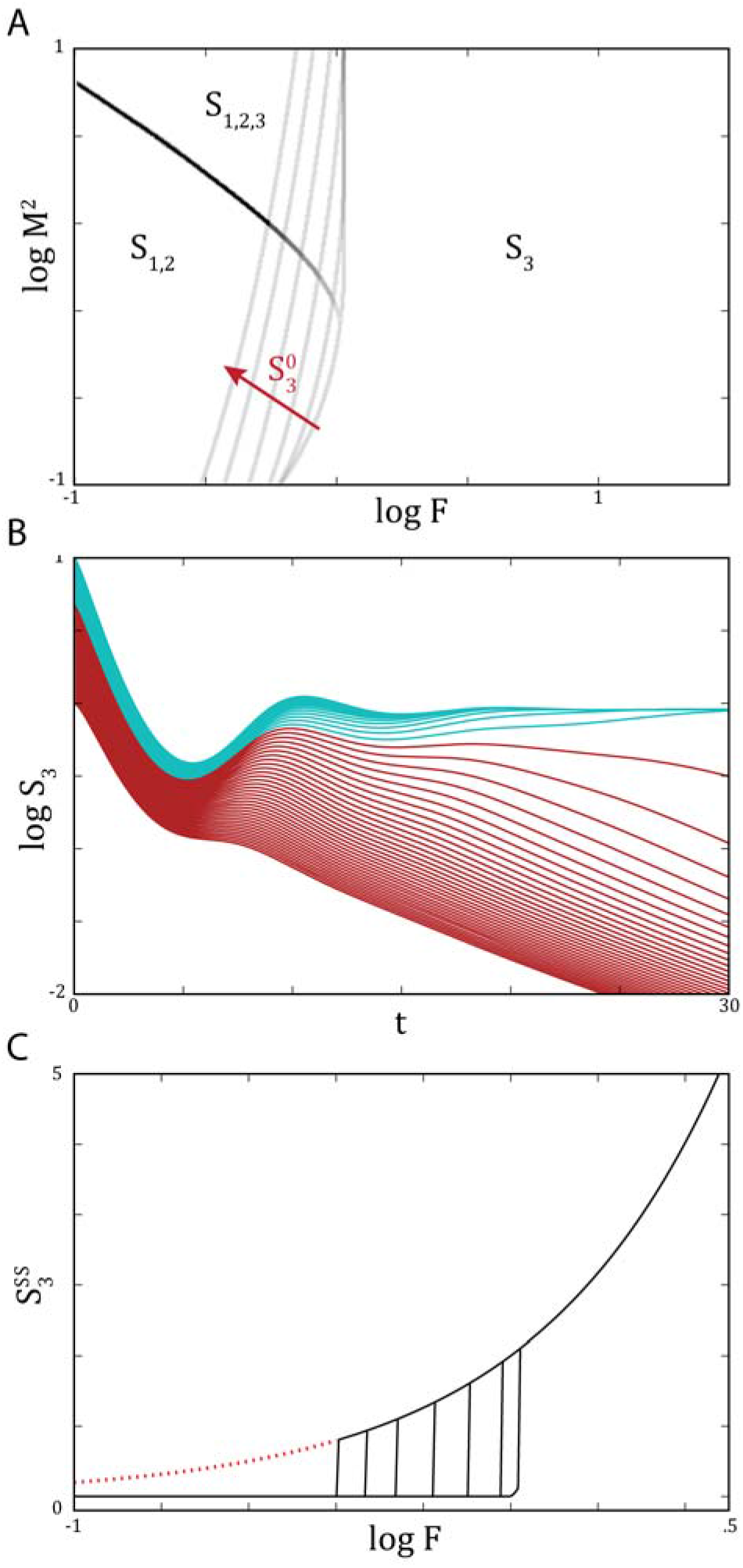
Initial invader density leads to dichotomy in mixed strategy invasion. (A) Phase i diagram of mixed strategy invasion. Shifted boundaries represent stepwise 10 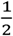 times increases in initial invader density (10^0^∼10^3^, along the arrow). Intersections of phase boundaries specify triple critical points. (B) Time trajectories across initial invader densities for successful (cyan) and unsuccessful (red) invasions at *M* = 1, *F* = 0.9. Bifurcation happens at 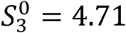. (C) Numerical steady state invader abundance along *M* = 0. Shifted transitions define bistable regime for initial invader densities 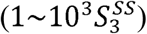 with 10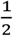 times stepwise increases. Invader-only steady state indicated by the red dashed line. This regime is theoretically possible but requires unrealistically high initial invader density. Non-varied parameters and initial conditions of native system are set as 1. Production rates of metabolite maintain the relation *β*_1_ = *β*_2_. Fitness defined as *F* = (*µ*_3_ + *µ*_3,2_)/*δ*_3_, with *µ*_3,1_ = *µ*_3,2_ and varied by changing *δ*_3_.

The existence of the triple point reveals an important tradeoff between competitive and parasitic strategies. Although competitive invasion is more aggressive, it is thwarted by private metabolite exchange amongst the native community. The mixed strategy invader effectively counters this defense by suppressing metabolite levels with parasitic growth, which opens up the niche for competitive growth. To be effective with this strategy, the invader must be abundant enough to rapidly consume low concentrations of metabolite and disrupt the native community trading network. We therefore investigated the effect of initial invader abundance on the phase boundaries of final community composition in mixed strategy invasion. We found that increasing initial invader abundance shifts the boundary defining replacement of the native community (Fig. 4A). Communities near the phase boundaries, particularly around the triple point, have the highest sensitivity to initial conditions. In these regimes, minute changes to invader abundance induce diverging trajectories to distinct stable states, i.e., invader dominance versus extinction (Fig. 4B). Theoretically, increase in invader density can continuously push this boundary, but requires invader densities orders of magnitude greater than the steady state abundance (Fig. 4C). Physical constraints in a real system therefore confine this bistability to regions nearest the phase boundaries. This result suggests that experimentally observed density dependence in community assembly may be driven by invader induced environmental remodeling around the triple point through resource consumption [7,9]. Our deterministic model predicts the conditions under which tipping points emerge and stochasticity can dictate the system dynamics [7,9].

### 2.4 Environmental resource

The behavior of mixed strategy invaders suggests that resource availability can modulate phase transitions in community composition. To generalize this concept, we added an external source to one metabolite (Fig. 5A). To prevent continuous buildup of resources after species extinction, we also included small dilution terms. With these modifications we evaluated steady state composition of the cross-feeding community under competitive invasion. We focus on this case as it allows us to individually drive one of the two native species and break symmetry.

**Figure 5.**
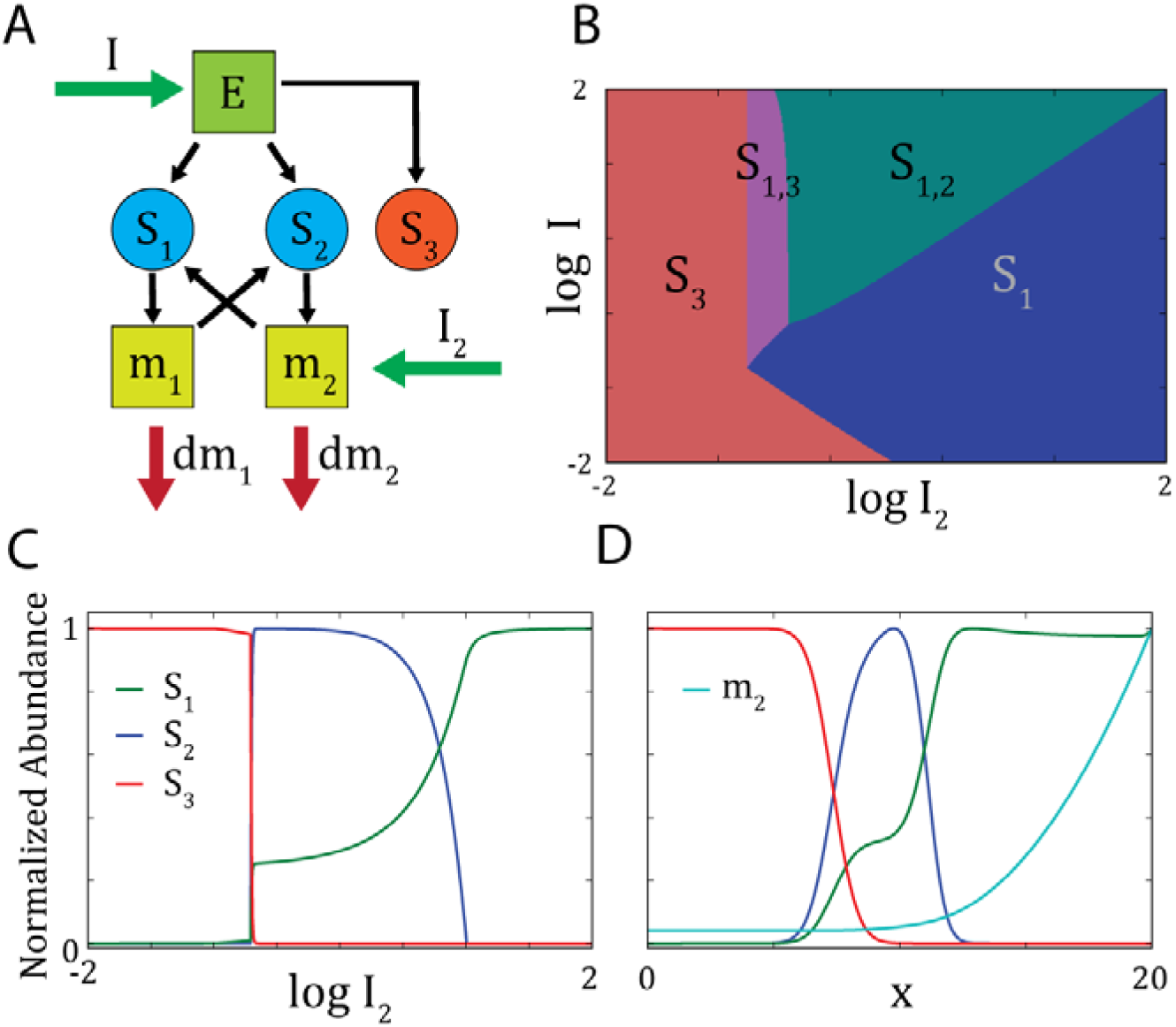
Environmental resource availability dictates invasion outcomes. (A) Community structure under competitive invasion with feeds *I* and *I*_2_ to energy and *m*_2_ respectively. Metabolites are diluted at the rate *d* = 0.1. (B) Phase diagram of steady state community composition across environmental feed rates. Coexisting species are specified for each phase. (C) Species abundance plotted against *I*_2_ when *I* = 10. (D) Spatial distribution of species abundance directed by diffusion of metabolite from a point source. Energy is uniformly fed at a rate *I* = 10; metabolite is fed at the right boundary, *I*_2_(*x* = 20) = 100. Diffusion coefficient of each resource is set as 1. Species diffusion coefficient is 0.001 to account for cell mixing. Other non-varied parameters are all set as 1.

For numerical solutions, we fixed species variables and generated a phase diagram of community composition across feed rates of energy and metabolite, *m*_2_ (Fig. 5B). With increasing feed of the metabolite, the system is driven through several fixed points: invader-only, coexistence of invader and fed species, native community only, and finally fed species only. The transition between invader and fed species coexistence to the native community only phase is first order, and the other transitions are second order (Fig. 5C).

The resource-induced phase transitions suggest continuous spatial gradients of external resources can establish sharp boundaries between regions that diverge in species composition. To implement this idea, we constructed a metabolite gradient through diffusive flux from a continuous point source in a one-dimensional system using numerical simulations (see Methods). We observed nonlinear transitions in species spatial distributions, specifically, a banded pattern as predicted from the phase diagram (Fig. 5D). In this case, the metabolite serves as a ‘morphogen’ that induces spatial pattern formation through species interactions.

## 3. Discussion

In natural ecosystems, community composition often undergoes dramatic transitions in response to habitat changes and invasion of new species [2,6]. Here, we model these phenomena in a mutualistic cross-feeding community to solve for system behaviors during these transitions. The key idea to make this model analytically tractable is that transitions between fully developed communities occur within the manifold of constrained resource abundance as species compete to fill the carrying capacity of the system. With this treatment, we observe rich phase transitions in the system as an emergent property of the interactions in the community network, which can be described using two simple system variables, the mutualism of native species and the fitness of invading species. These phase transitions appear to be insensitive to specific details of the model. For example, while we focus on the analytical results of linear growth in the current work, similar transition behaviors are also observed in numerical simulations using saturating growth and production rates.

At the phase boundaries, the discontinuity in invader abundance specifies the thresholds in the invader fitness and density for community entry. This forms an effective barrier as a collective property of the native community and presents greater resistance to invasion than either single species acting alone. As the emergence of this barrier is due exclusively to generic ecological features such as the extent of niche overlap, we expect these transitions to be ubiquitous in ecological systems. This result provides a mechanistic insight in explaining the difficulty in introducing new species to mutualistic ecosystems (e.g., probiotic therapies that aim to introduce new microbial species to change commensal microbiota), even when the new species can outcompete every native species separately [8].

The discontinuity also suggests that small changes in system parameters can induce abrupt and seemly stochastic changes in the community composition due to high susceptibility near the phase boundaries. The order of the discontinuity governs how the changes may initiate, which in turn provides distinct early signals to forecast the development of such changes in ecosystems [30–33]. Specifically, the effective energy barrier between alternative states during discontinuous first-order transitions dictates that the rate-limiting step is the slow nucleation of new states, which are typically initially localized in space and then rapidly spread across a habitat [16]. In this scenario, the nucleation events are useful early signals in predicting critical system-wide changes. Reversely, ecosystem restoration may be also achieved through local nucleation of healthy habitat [34]. In contrast, the lack of energy barrier during continuous second-order transitions prohibits the stable coexistence of two states in space, given uniform environmental conditions throughout the whole ecosystem [16]. Instead, systems at second-order phase transitions are expected to exhibit large, diverging fluctuations in time [16], which have been previously observed in ecological systems and interpreted as “critical slowing down” for early signal of ecological catastrophe [30]. While prior work in ecology often treats temporal and spatial fluctuations equivalently, our analyses indicate that the distinction between time and space is important for monitoring and managing ecosystems.

Another important feature of these phase behaviors is that they are dictated by resource constraints. This feature has several implications. First, the extent of overlap in resource requirements between native and invading species determines both the order of phase transitions and the location of phase boundaries. This fundamental property implies that similar phase transitions should also emerge in more complex ecosystems, which can contain a large number of species interacting through various mechanisms, as long as these species compete for overlapping resources. Second, we show that tuning resource availability through invader inoculation and external resource feeds can shift existing phase boundaries and create novel phases. This is because the strength of the interaction between native species depends on the abundance of private metabolites, which is in turn modulated by consumption and external input. Such variable interactions cannot be captured in pairwise models [14,15], and thereby highlight the necessity of modeling the resources explicitly. Finally, our results demonstrate the use of resources to control the outcome of invasions, and construct spatial, ecological patterns through resource gradients. The predictions made by our model suggest design principles and control mechanisms for ecological engineering. In particular, with the increasing accessibility to engineering complex biosynthetic interactions, our model provides a level of simplicity that can be fully implemented using obvious experimental systems such as syntrophic microbial strains with knockouts of one or more essential amino acid pathways [23,35,26]. The ability to drive these systems across multiple phase transitions using exogenously supplied amino acids provides a set of readily tractable controls orthogonal to traditional gene regulation techniques.

In summary, by analytically solving for species-resource interactions along a low-resource manifold, we investigated a mutualistic community under species invasion through several different invading mechanisms, ranging from competition to parasitism. We identified phase transitions that shift the system between several distinct stable states, including the persistence of mutualistic species, the coexistence of mutualists and invading species, and the replacement of mutualistic species by invaders. The order of discontinuity at these transitions offers insights to the underlying community network and suggests strategies to predict and control the outcomes of species invasion to ecosystems.

## 4. Methods

*Stability analyses*. For *P*_1_, we calculated the Jacobian and numerically identified the largest real eigenvalue component to be always negative for all parameter ranges. We then expanded the Jacobian to include competitive invader and calculated the largest eigenvalues along the parameter sweep shown in Fig. 2B. The largest eigenvalue of Pi flips sign to become positive at the phase boundary. Absence of native community at *P*_2_ prevents dynamic response to metabolite perturbation, resulting in a zero eigenvalue in all regimes. Numerical perturbations confirmed the attractor state of *P*_2_ at high fitness regimes. Similarly, the largest eigenvalues of both *P*_1_ and *P*_3_ switch sign at the phase boundary, confirming stability assertions.

*Numerical analyses*. Numerical solutions of ordinary differential equations were obtained through the odeint integrator function in SciPy 2017. Initial conditions were set to 1, unless noted otherwise, and species presence was thresholded at an abundance of 10^-3^. Spatial simulations were performed using matrix diffusion in a one-dimensional model with no flux boundary conditions. Species and resources were initialized with uniform conditions, and asymmetry was introduced through a constant flux of the metabolite in boundary cells on one end of the system.

## Acknowledgement

We thank S. Wan for technical assistance, and Jian Qin and Will Ludington for stimulating discussions and critical reading of the manuscript. This work is funded by Volkswagen Foundation (No. 94819). S.B. is supported by a NIH CMB training grant (T32GM007276). B.W. is supported by a CASI award from the Burroughs Wellcome Fund.

